# Nose-to-Knee Transplantation for Osteochondral Regeneration: A Proof-of-Concept Study in Sheep Employing Ceramic-Hydrogel Osteochondral Grafts

**DOI:** 10.64898/2026.10.09.757945

**Authors:** Philipp Fisch, Iman Roohani, Simone Ponta, Siyi Chen, Nathan Alibert, David Fercher, Karina Klein, Brigitte von Rechenberg, Marcy Zenobi-Wong, Salim Darwiche, Hala Zreiqat, Anna Puiggali-Jou

## Abstract

Osteochondral defects remain a major clinical challenge, frequently progressing to osteoarthritis and ultimately requiring joint replacement. Despite advances in regenerative strategies, current approaches often fail to restore the structural and functional complexity of the osteochondral unit, resulting in poor integration and limited long-term performance. Here, we developed autologous osteochondral grafts combining mechanically mature nasal chondrocyte-derived cartilage with a 3D-printed strontium–hardystonite–gahnite trabecular bone scaffold, which was tested in an osteochondral defect model in sheep. Nasal chondrocytes were encapsulated in a hyaluronan transglutaminase-alginate hydrogel generating hyaline-like cartilage with physiologically relevant compressive properties (1.0 ± 0.2 MPa) after 6 weeks of in vitro maturation. µCT scans of the osteochondral unit were used to design biomimetic trabecular bone scaffolds. Controlled infiltration of the hydrogel into the trabecular pores and subsequent cartilage maturation created a mechanical interlock between cartilage and bone layer, reaching an interfacial shear strength of 201 ± 42 kPa. The translational potential of this approach was evaluated in a proof-of-concept ovine study. The engineered cartilage remained attached to the bone scaffold for three months following implantation, but osteochondral repair was limited by implant sinking, likely caused by inadequate implant fixation in the defect, resulting in micromotion, synovial fluid infiltration, osteolysis and subsequent bone cyst formation. Together, this work demonstrates the feasibility of engineering mechanically mature autologous osteochondral grafts while identifying subchondral bone integration as the primary bottleneck limiting successful osteochondral regeneration and future clinical translation.

## Introduction

The osteochondral (OC) unit is a hierarchical structure composed of hyaline cartilage and subchondral bone that provides load distribution and joint stability.^1,2^ Although these tissues are adjacent, they exhibit distinct oxygenation, vascularization, and mechanical properties, making the engineering of the OC unit challenging.^3^ Traumatic OC injuries are common, particularly in sports-related activities, and frequently result in pain and impaired joint function.^4^ Despite existing clinical interventions^5^, OC lesions remain a major risk factor for progression to osteoarthritis (OA) and eventual joint arthroplasty.^6^ OA is a leading cause of global disability and represents a growing socioeconomic burden within ageing populations.^7^ In parallel, the demand for joint replacement surgery continues to increase. In Switzerland, the Swiss Implant Registry (SIRIS) reported 24,140 primary knee arthroplasties in 2023, compared to 18,546 in 2019, with a 2-year revision rate of 3.4% (95% CI 3.3–3.6%).^8^ These data highlight the need for regenerative strategies capable of restoring OC tissue and delaying progression to joint replacement.

Current clinical strategies for OC repair include autologous and allogeneic OC grafts (OCG). While these treatments partially preserve native tissue architecture and function, their use is limited by donor-site morbidity, restricted graft availability, and concerns regarding tissue supply and disease transmission.^9,10^ To overcome these limitations, first-generation acellular OC scaffolds, including TruFit (Smith & Nephew, USA)^11,12^, ChondroMimetic (Collagen Solutions, UK)^13^, MaioRegen (Fin-Ceramica S.p.A., Italy)^14^, and Agili-C (CartiHeal Ltd, Israel)^15^, were developed as off-the-shelf alternatives. As cell-free constructs, these scaffolds rely on the host response and therefore show variable clinical outcomes. TruFit was withdrawn from the market in 2013 due to inconsistent cartilage repair and insufficient subchondral bone regeneration.^12,16,17^ ChondroMimetic demonstrated partial structural repair in preclinical large-animal studies and early clinical investigations, but has not achieved widespread clinical adoption.^13,18^ Newer scaffolds, including MaioRegen^14,19,20^ and Agili-C^21,22^, have shown improved short- and mid-term outcomes, although limitations persist, including incomplete tissue integration and uncertain long-term durability. These heterogeneous outcomes are largely attributed to the intrinsic repair mechanism of acellular scaffolds, which depends on endogenous progenitor cell recruitment and differentiation within a biologically variable and inflammatory environment, frequently resulting in incomplete regeneration of cartilage and subchondral bone.^23–25^

These limitations highlight the need for next-generation cellular OC grafts capable of promoting subchondral bone regeneration while providing a mechanically stable cartilage layer that integrates with its surrounding tissue and sustains joint loading. Multiple research groups have pursued this strategy using biphasic and multiphasic constructs, including ceramic-printed bone scaffolds combined with melt-electrowritten reinforced cartilage layers^26^ and metal^27^ or polycaprolactone^28^ (PCL)-printed bone scaffolds integrated with cartilage microtissues. de Ruijter et al. demonstrated that the mechanical stability of the cartilage phase is a critical determinant of OC graft performance.^29^ However, even advanced multiphasic scaffolds designed to replicate native OC gradients remain susceptible to interfacial separation and inadequate cell retention under physiological loading conditions, thereby limiting long-term graft stability and function.^30–32^ We therefore hypothesized that a mechanically mature cartilage graft with sufficient intrinsic stiffness would establish a stable physical interlock with a porous trabecular ceramic scaffold, thereby maintaining structural integrity under joint loading.

Cell sources for OC cartilage tissue engineering commonly include articular chondrocytes, osteoblasts, synoviocytes, and mesenchymal stem cells (MSCs). Articular chondrocytes have been used clinically for decades in cartilage repair approaches such as autologous chondrocyte implantation (ACI)^33–35^ and matrix-assisted autologous chondrocyte implantation (MACI). ^36,37^ Their application is supported by their ability to synthesize cartilage-specific extracellular matrix (ECM).^38,39^ However, clinical application is limited by donor-site morbidity, low cell yield, and dedifferentiation during in vitro expansion, which can compromise stable hyaline cartilage formation. MSCs derived from bone marrow^40^, adipose tissue^41^, and synovium^42^ have emerged as alternative cell sources due to their accessibility, proliferative capacity, and osteochondrogenic potential. However, hypertrophy and ectopic calcification remain important limitations for stable cartilage regeneration. Nasal chondrocytes, which originate from the neural crest, have therefore gained increasing attention due to their low harvest morbidity, high proliferative capacity, and superior chondrogenic stability under inflammatory conditions compared to articular chondrocytes.^43–45^ However, a recent clinical study has highlighted the need for appropriate ex vivo maturation, where prolonged ex vivo maturation was associated with improved clinical outcomes.^45^ We recently demonstrated near native tissue maturation using auricular chondrocytes encapsulated in a hyaluronan transglutaminase-alginate (HATG-Alg) bioink.^46^ As nasal chondrocytes originate from the same neural crest origin, we hypothesized that they would possess a similar intrinsic cartilage regeneration capacity as auricular chondrocytes when cultured under these conditions.

In the present study, we leverage the HATG-Alg system and developed a “nose-to-knee” strategy for OC regeneration by integrating HATG-Alg cartilage grafts^46–50^ with a 3D-printed strontium-hardystonite-gahnite (Sr-HT-Gahnite) bone ceramic exhibiting osteoconductive properties.^51,52^ The porous ceramic architecture was designed using µCT-derived sheep trabecular bone structures, enabling controlled infiltration of the HATG-Alg hydrogel and formation of a mechanically interlocked transition zone between cartilage and bone. Autologous nasal chondrocytes embedded within the hydrogel generated cartilage tissue with hyaline-like characteristics, while the ceramic phase provided structural support for subchondral bone regeneration. We hypothesized that combining a mechanically mature cartilage graft with an anatomically optimized ceramic scaffold would promote functional OC regeneration in vivo. As proof of concept, grafts were matured in vitro and implanted into OC defects in the medial and lateral femoral condyles in sheep stifle joints. Tissue repair was evaluated after 3 months using macroscopic scoring, µCT, and histological analyses (Figure 1). While the engineered cartilage remained integrated with the ceramic scaffold and was retained after implantation, subchondral bone integration emerged as the primary factor limiting successful OC repair in the load-bearing ovine model.

**Figure 1.**
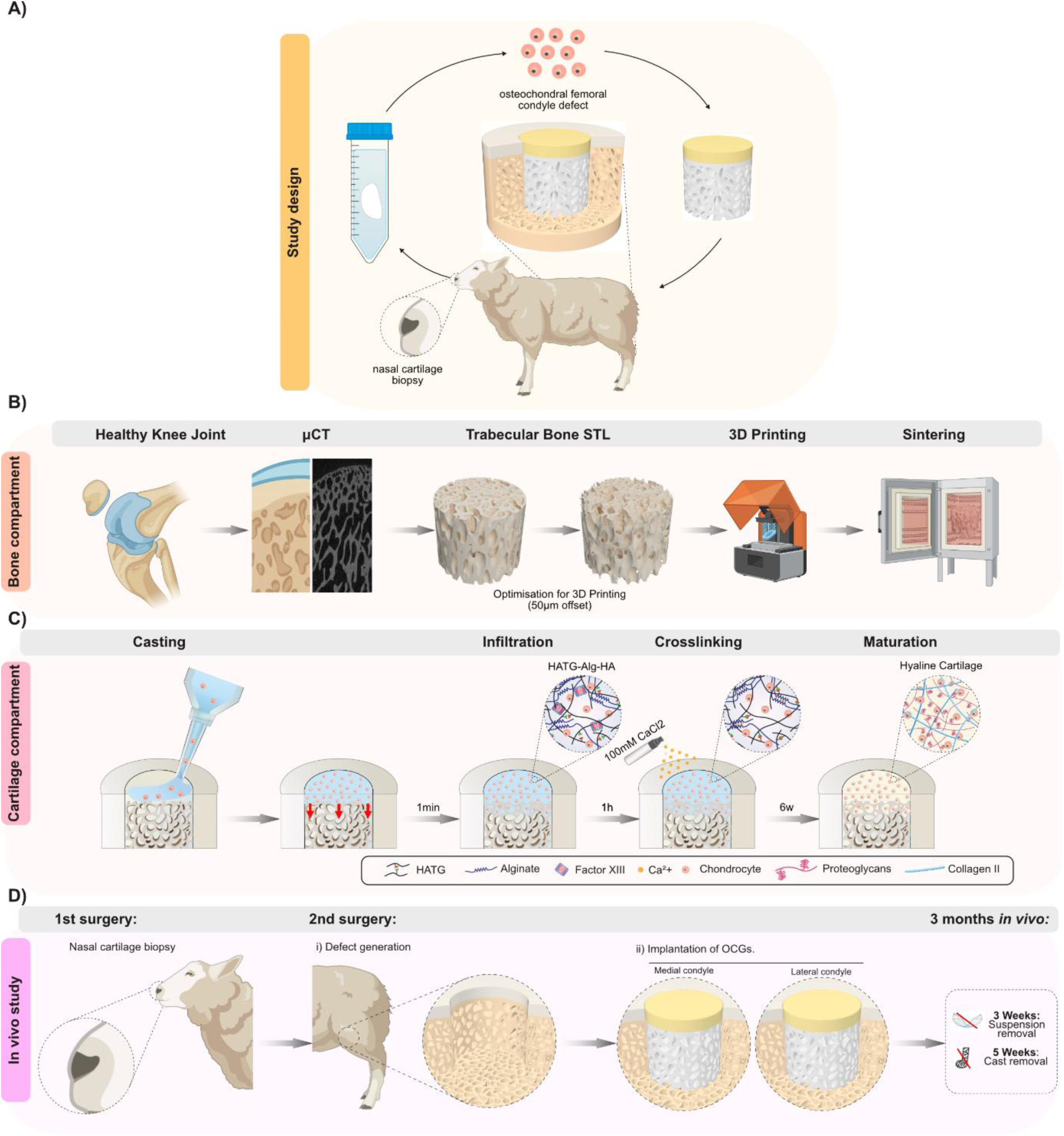
Schematic representation of the experimental approach. (**A**) Autologous “nose-to-knee” transplantation approach. A nasal cartilage biopsy was collected, nasal chondrocytes isolated, and in combination with a hyaluronan-based hydrogel, casted on a ceramic bone scaffold to create an autologous osteochondral graft (Adapted from BioRender. P. Jou, A. (2027) https://BioRender.com/xruh06y). After maturation, the osteochondral graft was implanted in osteochondral defects in the stifle joint of the same sheep. (**B**) Design and manufacturing of the ceramic bone scaffolds. An osteochondral plug of a sheep stifle joint (Adapted from BioRender. P. Jou, A. (2027) https://BioRender.com/xruh06y) was imaged using µCT and the trabecular bone isolated. The resulting trabecular bone graft was 3D printed using a ceramic-resin and subsequently sintered. (**C**) Sheep nasal chondrocytes were combined with an HATG-Alg-HA hydrogel, cased on top of the ceramic bone scaffold and allowed to infiltrate the trabecular pores (red arrows). After infiltration, the hydrogel was crosslinked with CaCl2 and matured for 6 weeks to allow hyaline cartilage to form. (**D**) Detailed “nose-to-knee” transplantation (Adapted from BioRender. P. Jou, A. (2027) https://BioRender.com/xruh06y). In a 1^st^ surgery, nasal cartilage was collected and the osteochondral grafts generated as depicted in (C). Two osteochondral defects were generated in the medial and lateral condyle of the sheep’s stifle joint. For the first 3 weeks, sheep were kept in suspension, after 5 weeks, the cast was removed, and the total in vivo period spanned 12 weeks.

## Results and discussion

### Design and fabrication of trabecular bone graft using ceramic 3D printing

The subchondral bone layer is commonly fabricated from ceramics such as hydroxyapatite (HAp) and tricalcium phosphate (TCP), but their mechanical performance is limited when porosity levels are designed to match trabecular bone. To overcome these limitations, we selected a strontium–hardystonite–gahnite (Sr-HT-Gahnite) ceramic, which has been shown to enhance mechanical strength and bioactivity.^53^ We generated 3D models from µCT images of the OC unit of sheep cadaver joints to mimic the native trabecular bone architecture (Figure 1B). The trabecular region below the subchondral bone plate was selected, thresholded, converted into a 3D model, and printed using an Sr-HT-Gahnite ceramic resin on a custom digital light processing (DLP) printer.^54^ Once the prints were sintered and sterilized, we cast our hydrogel onto them to replicate the OC unit.

Post-printing, the trabecular bone ceramics were scanned and reconstructed from µCT images to analyze trabecular bone specific metrics and compare them to sheep trabecular bone. From the images, we observed matching features between the 3D model and the printed ceramics (Figure 2A,B, trabecular bone graft model – blue arrows). However, in areas where small pores dominated, the ceramic resin was difficult to remove, resulting in fused trabeculae (red arrows). We therefore used the surface with the highest fraction of open pores to cast our cartilage hydrogel. To compare the 3D printed bone scaffolds to trabecular bone samples from the medial and lateral condyles of healthy sheep joints, we analyzed several µCT-specific trabecular parameters and compared them (Figure 2B, structural comparison). Bone ceramics showed a higher bone volume percentage and lower trabecular thickness and separation. While some of these differences were significant, the printed bone ceramics generally fell within the same order of magnitude of these features with a bone volume percentage of 48.9 ± 3.3% (native medial: 43.0 ± 6.6 %, native lateral: 29.7 ± 5.8 %), a trabecular separation of 345.2 ± 24.1 µm (native medial: 429.3 ± 58.2 µm, native lateral: 527.9 ± 59.2 µm), and a trabecular thickness of 102.0 ± 5.1 µm (native medial: 204.0 ± 18.2 µm, native lateral: 162.6 ± 16.6 µm). SEM images revealed microscopic pores and surface roughness on top of the printed ceramic, ideal for cell adhesion (Figure 2B, SEM assessment).^55^ Overall, these results indicate that the ceramic printing process closely mimicked the trabecular structure of native sheep trabecular bone.

**Figure 2.**
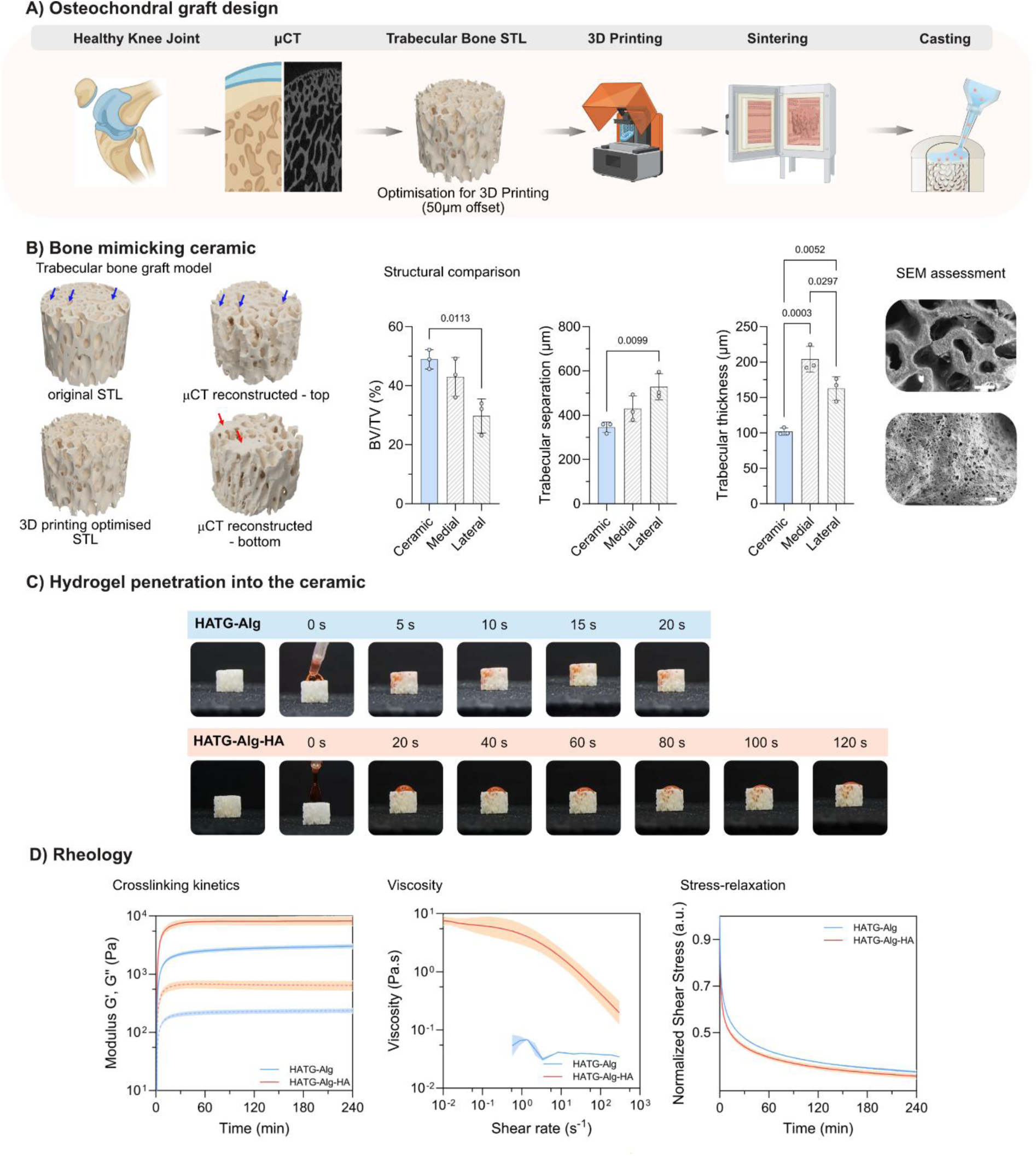
Bioinspired fabrication and characterization of a biphasic osteochondral graft. (**A**) Procedure for the fabrication of the osteochondral graft: A µCT scan of a health sheep joint was taken, segmented and 3D reconstructed to generate a 3D model of the trabecular structure (Adapted from BioRender. P. Jou, A. (2027) https://BioRender.com/xruh06y). The 3D model was optimized for printing and 3D-printed on a high-resolution DLP printer using a Sr-HT-Gahnite ceramic. Post-printing, the bone scaffold was sintered and the cartilage hydrogel cast onto it. (**B**) Structural characterization of the ceramic scaffold. Trabecular bone graft model: the printed ceramic scaffolds were scanned with µCT, reconstructed, and compared with the original 3D model. Blue arrows indicate matching trabecular features between the 3D model and the printed bone scaffold, while red arrows indicate closed trabecular pores. Structural comparison: bone volume (BV)/ total volume (TV)%, trabecular separation and trabecular thickness as analyzed from µCT scans. SEM assessment of bone ceramic. Scale bar = 400 µm; inset = 20 µm. (**C**) Cartilage hydrogel precursors infiltration into the ceramic comparing HATG-Alg and HATG-Alg-HA. (**D**) Rheological characterization for crosslinking kinetics (G’: full line, G’’: dotted line), viscosity and stress relaxation of HATG-Alg and HATG-Alg-HA.

We next investigated casting our cartilage-based hydrogel on top of the ceramic scaffold. To achieve controlled interpenetration of our hydrogel pre-courser into the ceramic scaffold, we developed two hydrogels based on our previously reported HATG-Alg bioink with varying viscosities. The first hydrogel consisting of 0.5% HATG and 0.25% Alg (HATG-Alg) displayed a negligible viscosity and rapidly infiltrated the entire ceramic construct within 5 seconds (Figure 2C), making crosslinking of the hydrogel before complete infiltration into the ceramic scaffold impossible. To circumvent this, we increased the hydrogel’s viscosity by incorporating unmodified hyaluronan (HATG-Alg-HA). The addition of HA to the hydrogel substantially increased its viscosity and led to a slower infiltration of the bone ceramic, with a portion of the hydrogel remaining on top of the ceramic after 60 seconds of casting (Figure 2D, viscosity). This provided sufficient time for calcium-triggered crosslinking of HATG and ionic crosslinking of Alg,^47^ thereby preventing complete hydrogel infiltration. The addition of HA further increased the hydrogels storage (HATG-Alg: 3.0 ± 0.3 kPa, HATG-Alg-HA: 8.3 ± 1.6 kPa) and loss (HATG-Alg: 0.2 ± 0.03 kPa, HATG-Alg-HA: 0.6 ± 0.13 kPa) modulus upon crosslinking (Figure 2D, crosslinking kinetics), and had a small but significant effect on its stress-relaxing properties related to the free, unmodified HA chains (HATG-Alg: 262 ± 26 s, HATG-Alg-HA: 833 ± 103 s, Figure 2D, stress-relaxation).

### Nasal chondrocytes regenerate functionally mature hyaline cartilage

The stability of OCG and their integration with the host tissue strongly depends on the maturation of cartilage. After casting and crosslinking, our biomaterial reaches 8.3 ± 1.6 kPa in storage modulus, insufficient to sustain the loading experienced once implanted in the joint. We therefore evaluated four different chondrogenic cell types – synoviocytes, articular, auricular, and nasal chondrocytes in HATG-Alg – for their chondrogenic regeneration potential.

All cell types remained viable for 3 weeks and exhibited second harmonic generation (SHG) signal, indicating collagen deposition (Figure S1). After 6 weeks of culture, varying degrees of cartilage regeneration could be identified based on biomechanical and histological readouts. Auricular chondrocytes reached the highest compressive modulus (692 ± 194 kPa), followed by nasal chondrocytes (459 ± 131 kPa), while articular chondrocytes and synoviocytes reached the lowest levels with 195 ± 54 kPa and 215 ± 13 kPa, respectively. Collagen I, a marker for fibrocartilage, was absent throughout nasal and auricular grafts but was expressed in articular and synoviocytes grafts. In alignment with the biomechanical properties, glycosaminoglycans (GAGs) and collagen II deposition was most pronounced in auricular and nasal grafts, while articular and synoviocytes grafts showed only minor deposition of GAGs and collagen II. Taken together, these data underline the potential of nasal and auricular chondrocytes as regenerative cell source for autologous cartilage grafts. However, as auricular chondrocytes are of elastic cartilage origin, characterized by a high degree of elastin in its extracellular matrix, they are less suitable for applications in the joint.^56^ We therefore proceeded with nasal chondrocytes as cell source for hyaline cartilage tissue repair, following recent clinical trials applying a nose-to-knee approach^43,45,57^.

To assess the long-term regenerative potential, we then combined nasal chondrocytes with HATG-Alg-HA and cultured 4 mm cartilage discs for up to 4 months (Figure 3A). Nasal chondrocytes remained viable (>80%) for up to 6 weeks and grafts showed the presence of collagen based on SHG signal (Figure 3B). After 6 weeks and 4 months, we compared the histological and biomechanical properties of these grafts to native sheep nasal (sNAS) and articular (sART) cartilage. Grafts showed the deposition of GAGs and collagen II, while collagen I remained low (Figure 3C). Although GAGs did not reach the same intensity as in sNAS and sART controls, showing a slight decline in intensity at 4 months compared to 6 weeks, our results underline the maturation of grafts into hyaline cartilage. Collagen X and alizarin red, markers for hypertrophy and calcification, were absent. Picrosirius red staining was distinct in sART cartilage compared to tissue engineered grafts and sNAS cartilage, potentially indicating differences in total collagen content between sNAS and sART. The mechanical properties followed the histological trends (Figure 3D). sNAS cartilage reached the highest instantaneous modulus (1753 ± 250 kPa), while grafts cultured for 6 weeks (1016 ± 169 kPa) and 4 months (805 ± 271 kPa), as well as sART cartilage (804 ± 163 kPa) reached similar moduli. More pronounced differences were observed in equilibrium modulus, with sNAS cartilage again reaching the highest values (719 ± 38 kPa), followed by sART cartilage (117 ± 79 kPa) and 6 weeks (52 ± 21 kPa) and 4 months (24 ± 23 kPa) old grafts. To further assess differences in ECM deposition, we performed biochemical quantification of total DNA and GAGs. The engineered grafts reached a dry weight of 8.8 ± 0.4% after 6 weeks and 7.7 ± 0.6% after 4 months, which remained markedly lower than that of native tissues (sART: 19.6 ± 2.2%; sNAS: 26.9 ± 1.1%; Figure 3Ei). DNA normalized to wet weight (%_WW_) indicated that the engineered grafts exhibited cellularity comparable to sNAS and more than twofold higher than sART (Figure 3Eii). GAG quantification followed the same trend observed histologically, with the highest GAG levels in sNAS cartilage and the lowest in grafts cultured for 4 months (Figure 3Eiii). However, when normalized to DNA, GAG per cell was highest in sART (Figure 3Eiv). In summary, we were able to engineer mature, mechanically competent hyaline cartilage, closely approximating the biomechanical and histological properties of sART cartilage using sNAS chondrocytes, underlining their suitability to engineer hyaline cartilage for OC applications in the joint.

**Figure 3.**
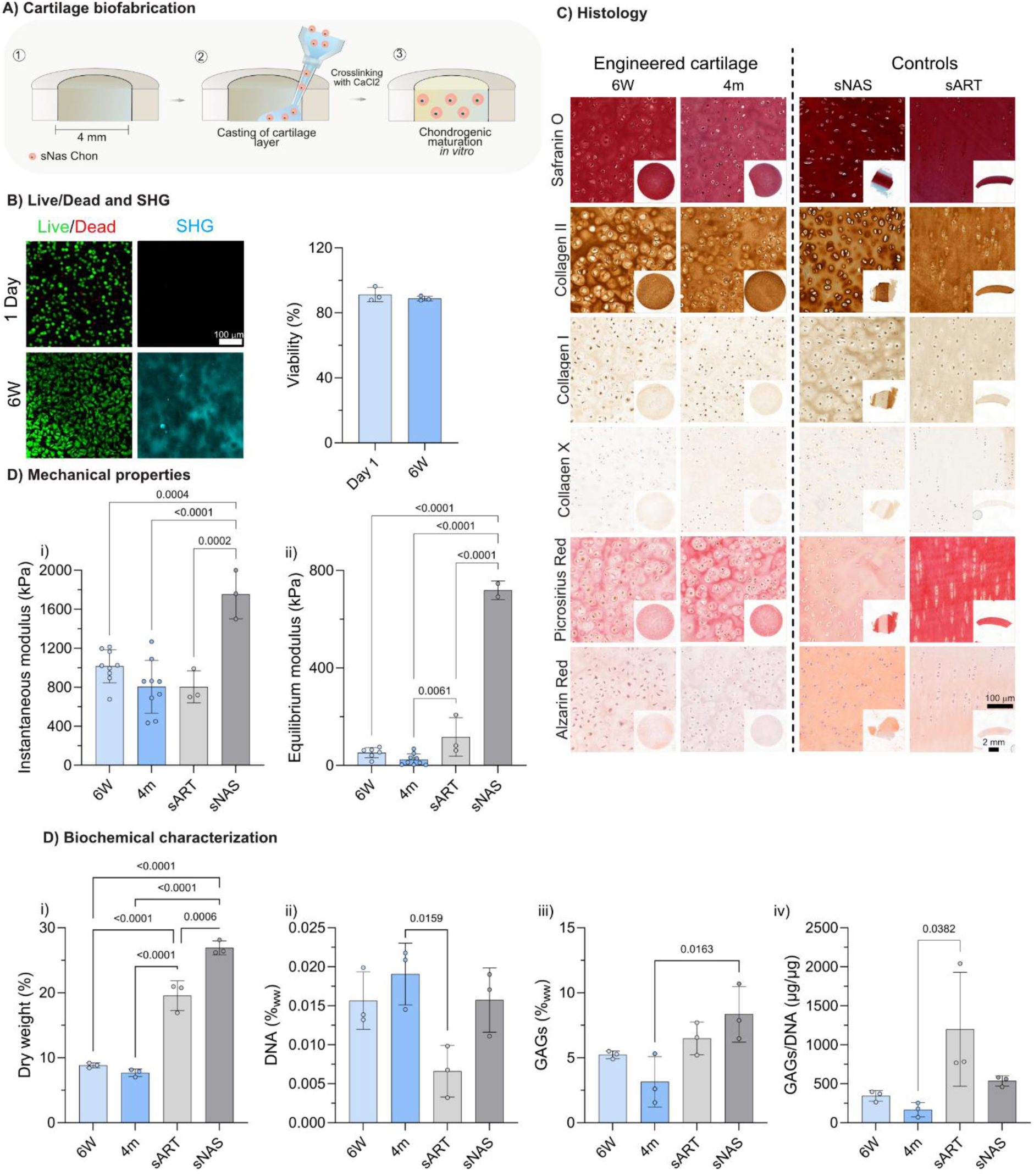
*In vitro* maturation and characterization of nasal chondrocyte–derived cartilage grafts. (A) Schematic of the fabrication process: (1) 4 mm molds preparation, (2) nasal chondrocytes encapsulation in a viscosity-enhanced hydrogel and casting into the molds, (3) constructs maturation in chondrogenic media for up to 4 months (4m). (B) Cell viability and matrix formation assessed by live/dead staining at day 1 and 6 weeks (6W), and collagen deposition visualized by second harmonic generation (SHG) imaging. (C) Histological and immunohistochemical characterization of engineered grafts (6W and 4m) compared to native sheep nasal cartilage (sNAS) and articular cartilage (sART), including Safranin O staining, immunostaining for collagen I, II, and X, picrosirius red, and alizarin red staining. Scale bar = 2 mm; inset = 100 µm. (D) Mechanical characterization showing the instantaneous (i) and equilibrium (ii) modulus of the different groups. (E) Biochemical quantification of ECM, including dry weight % (i), DNA%ww (ii), glycosaminoglycans %ww (iii), and GAGs/DNA ratio (iv).

### Interfacial integration of cartilage and bone through hydrogel infiltration

A major challenge in biofabricating the OC unit is achieving stable integration between the cartilage and bone compartments, which possess markedly different structural and mechanical properties. To fabricate the hybrid ceramic–hydrogel graft, we placed the ceramic scaffolds in PDMS molds and cast our hydrogel precursor on top. After casting, the hydrogel precursor was allowed to infiltrate the pores of the ceramic scaffold for 1 min, after which crosslinking was initiated through calcium chloride (Figure 4A). Immediately after casting, the hydrogel layer easily detached upon shearing but became increasingly stiff and resistant to delamination with prolonged culture (Figure 4B).

**Figure 4.**
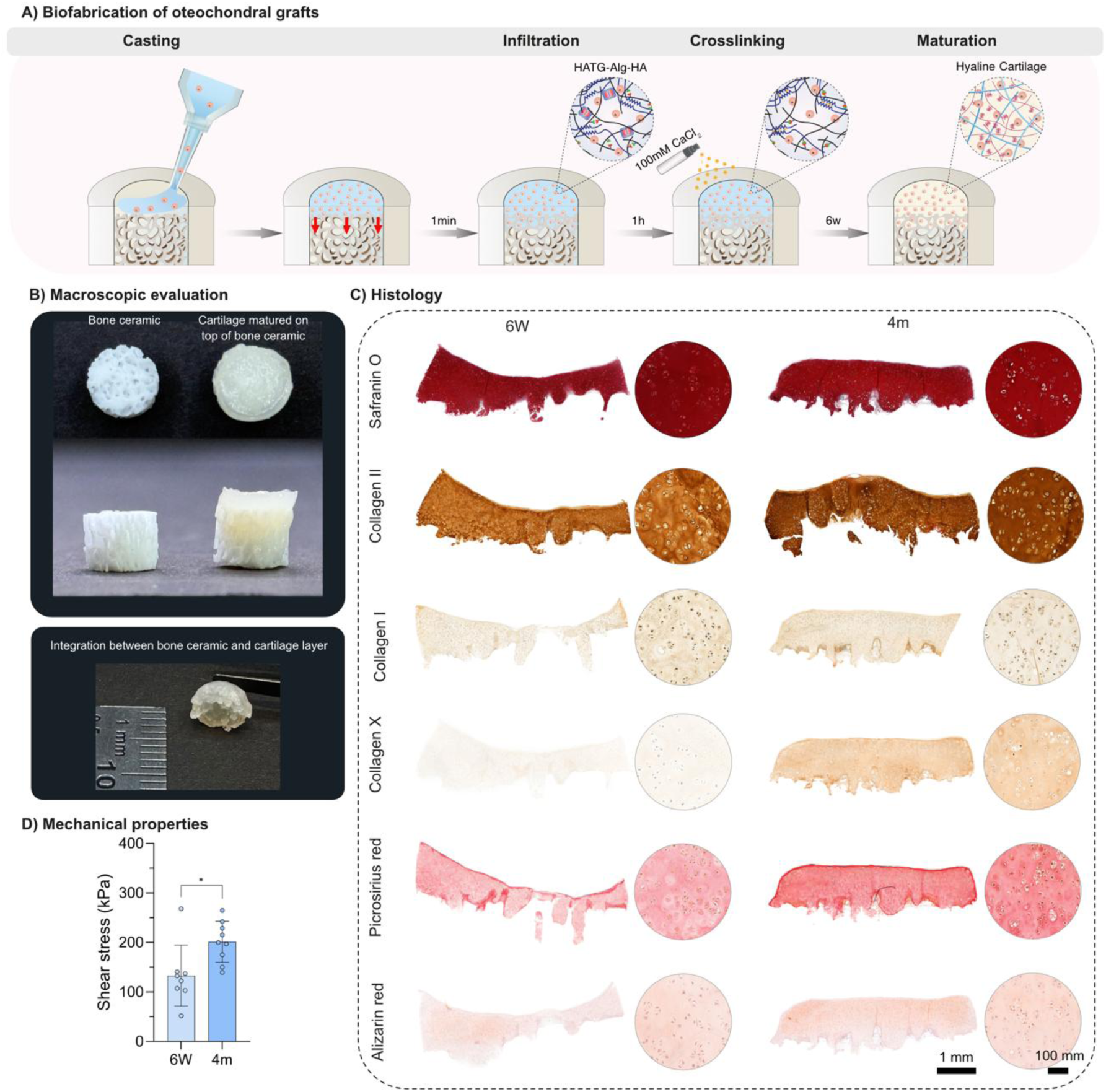
Biofabrication and in vitro maturation of osteochondral grafts (OCGs). (**A**) Schematic of the fabrication process: porous ceramic scaffolds were placed in PDMS molds, overlaid with a hydrogel containing sheep nasal chondrocytes, crosslinked with CaCl₂, and cultured for chondrogenic maturation. (**B**) Macroscopic appearance of OCGs showing the ceramic and engineered cartilage layers and their integration over time. (**C**) Histological analysis at 6 weeks (6W) and 4 months (4M) using Safranin O, collagen type II, I, and X, Picrosirius Red, and Alizarin Red. Scale bar = 1 mm; inset = 100 µm. (D) Interfacial shear testing showing the mechanical integration between cartilage and ceramic phases.

To evaluate bonding between cartilage and bone layer, the cartilage layer was sheared off the ceramic scaffold after 6 weeks and 4 months of culture and analyzed histologically (Figure 4C). As the engineered cartilage matures and stiffens, the interfacial bonding strength increases due to the stiffening of the hydrogel that infiltrates the porous ceramic structure. To quantify this integration, the shear stress required to detach the cartilage layer from the ceramic scaffold was measured. The detachment stress increased from 133 ± 61.6 kPa after 6 weeks to 201.4 ± 41.6 kPa after 4 months of culture (Figure 4D). These values approach the reported shear strength of the native cartilage–bone interface (0.5–1 MPa)^58^, indicating progressive interfacial reinforcement.

Furthermore, the ceramic scaffold did not impair cartilage maturation. The engineered tissue exhibited abundant glycosaminoglycans (GAGs) and collagen II deposition, while collagen I and collagen X, markers of fibro- and hypertrophic cartilage, remained low (Figure 4C). A modest increase in collagen X staining was observed after 4 months, suggesting the onset of mild hypertrophic changes during extended in vitro culture. Nevertheless, no mineralization was detected by Alizarin Red staining. Picrosirius Red staining further revealed a progressive increase in collagen deposition over time, with a stronger staining observed after 4 months of culture. Together, these results suggest that the engineered nasal sheep OCG develops sufficient interfacial strength to withstand mechanical loading and resist delamination.

### Proof-of-concept osteochondral ovine study

We next conducted a proof-of-concept study in an ovine model to evaluate the translational potential of this approach. The study was designed to assess the two-step surgical procedure, implant stability and integration as well as early regenerative outcomes at the 3-month timepoint in three sheep. During the first surgical procedure, a nasal cartilage biopsy (6 mm in diameter, ∼20 mg) was harvested from each animal, and nasal chondrocytes isolated, expanded, and used to generate OCGs, which were matured in vitro for 6 weeks. In a second surgical procedure, OC defects of 6.3 ± 0.1 mm in diameter and 6.4 ± 0.5 mm in height (with OCG implant dimensions matched accordingly) were created in the medial and lateral femoral condyles of one hind limb (Figure 5A). Each animal received OCGs containing its own engineered autologous cartilage.

**Figure 5.**
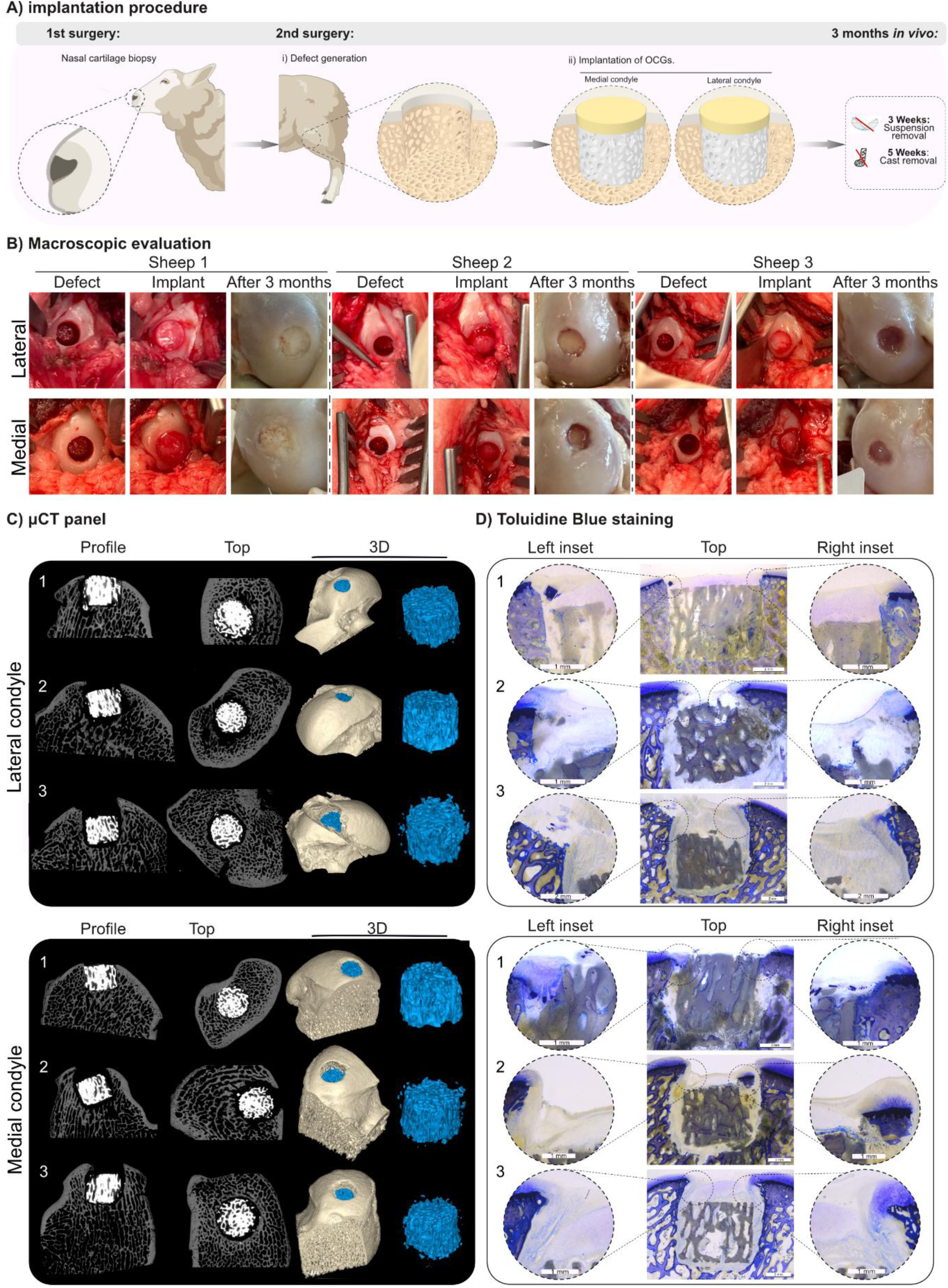
In vivo evaluation of osteochondral grafts (OCGs) in an ovine model. (**A**) Schematic representation of the two-step surgical procedure (Adapted from BioRender. P. Jou, A. (2027) https://BioRender.com/xruh06y). A nasal cartilage biopsy was harvested, and autologous chondrocytes were isolated, expanded, and used to generate engineered cartilage integrated into OCGs, which were matured in vitro for 6 weeks. In a second surgical intervention, osteochondral defects were created in the medial and lateral femoral condyles and filled with the corresponding OCGs. (**B**) Macroscopic evaluation of defect sites at implantation and after 3 months in vivo. (**C**) Micro-computed tomography (µCT) analysis of medial and lateral condyles, shown as profile, top, and 3D reconstructions. (D) Toluidine blue surface-staining of ground MMA sections highlighting cartilage (sulfated glycosaminoglycans) and mineralized bone. Scale bar = 2 mm; inset = 1 or 2 mm.

All animals tolerated the surgery well, showed no persistent lameness or abnormal behavior, and no clinical abnormalities were observed beyond those expected from the surgery. The insertion of the OCGs into the defect however posed difficult due to the irregular trabecular architecture of the bone graft which created an irregular surface and resulted in friction during insertion. The implants therefore had to be carefully positioned and aligned with the defect for insertion. Ultimately, all defects were successfully filled with OC grafts. Post-surgery, sheep were maintained in suspension for 3 weeks and the cast removed after 5 weeks. Some sheep did not accept a cast for the planned duration, prompting the veterinary team to remove the cast earlier than planned. This allowed for more stifle flexion for Sheep 1 after its time in the suspension nets. For sheep 2, the cast was removed one week prior to the planned timepoint, allowing flexion starting 4 weeks after surgery.

Three months post-surgery, repair at the implantation sites was compared with surrounding healthy tissue. Sheep 1 showed the best repair, with both OCGs remaining in place at the implantation site and cartilage covering parts of the construct (Figure 5B). In contrast, OCGs implanted in the medial and lateral condyle in sheep 2 and 3 sunk within the defect, although covered by cartilaginous tissue. Micro-computed tomography (µCT) of the defect sites further confirmed a tight fit between the OCG and bone with minimal osteolysis in sheep 1, which was restrained to the broken edge of the bone ceramic (Figure 5C) and may have occurred from wear particles activating the immune response, that in turn activates osteoclast activity.^59,60^ In contrast, the sunken samples of sheep 2 and 3 exhibited pronounced bone resorption around the implant in both the medial and lateral condyles, as indicated by the black areas between bone ceramic (white) and natural bone (grey). As the insertion of the implants was complicated by their irregular surface, a true press-fit could not be achieved, likely resulting in micromotion and progressive instability under repetitive joint loading leading to bone loss. This might have been aggravated by synovial fluid filling the gaps at the implant–bone interface, further promoting osteolysis.^61^

To evaluate tissue formation and integration further, we surface-stained 500-700 µm MMA sections with toluidine blue, enabling the visualization of sulfated GAGs within the cartilage region, appearing as purple, while mineralized bone matrix stained in shades of blue, with darker blue indicating newer bone formation (Figure 5D). Consistent with our previous observations, sheep 1 exhibited the best implant integration at the defect side and maintained its cartilage layer in the lateral side. We however also observed that the engineered cartilage layer reached below the tide mark due to the thicker engineered cartilage layer compared to the native cartilage layer. Sheep 2 and 3 on the other hand showed fibrous tissue surrounding the bone ceramic, creating a gap between native bone and implant. However, we were able to detect cartilage on 3 out of the 4 implants, indicating good integration between bone ceramic and engineered cartilage.

Lastly, 5 µm MMA thin sections were created. Due to the brittleness of the ceramic, the ceramic detached during the process. We were therefore only able to visualize the remaining engineered cartilage in greater detail (Figure 6). In 4 out of 6 cases, we were able to localize the engineered cartilage (marked with a white asterisk), which retained most of its GAGs, as evidenced by Safranin O and Toluidine Blue staining. In the remaining two cases, the section either did not contain the OCG (lateral, sheep 2) or parts of the section was lost in processing due to the challenges of sectioning these samples (medial, sheep 2), not allowing us to evaluate the cartilage. Furthermore, chondrocytes within the engineered cartilage retained their round morphology. Von Kossa Mac Neals staining was performed to visualize new bone formation. However, there was no indication of bone formation within the grafts, only fibrous tissue surrounding the implant, likely due to micromotions between the implant and the surrounding bone.

**Figure 6.**
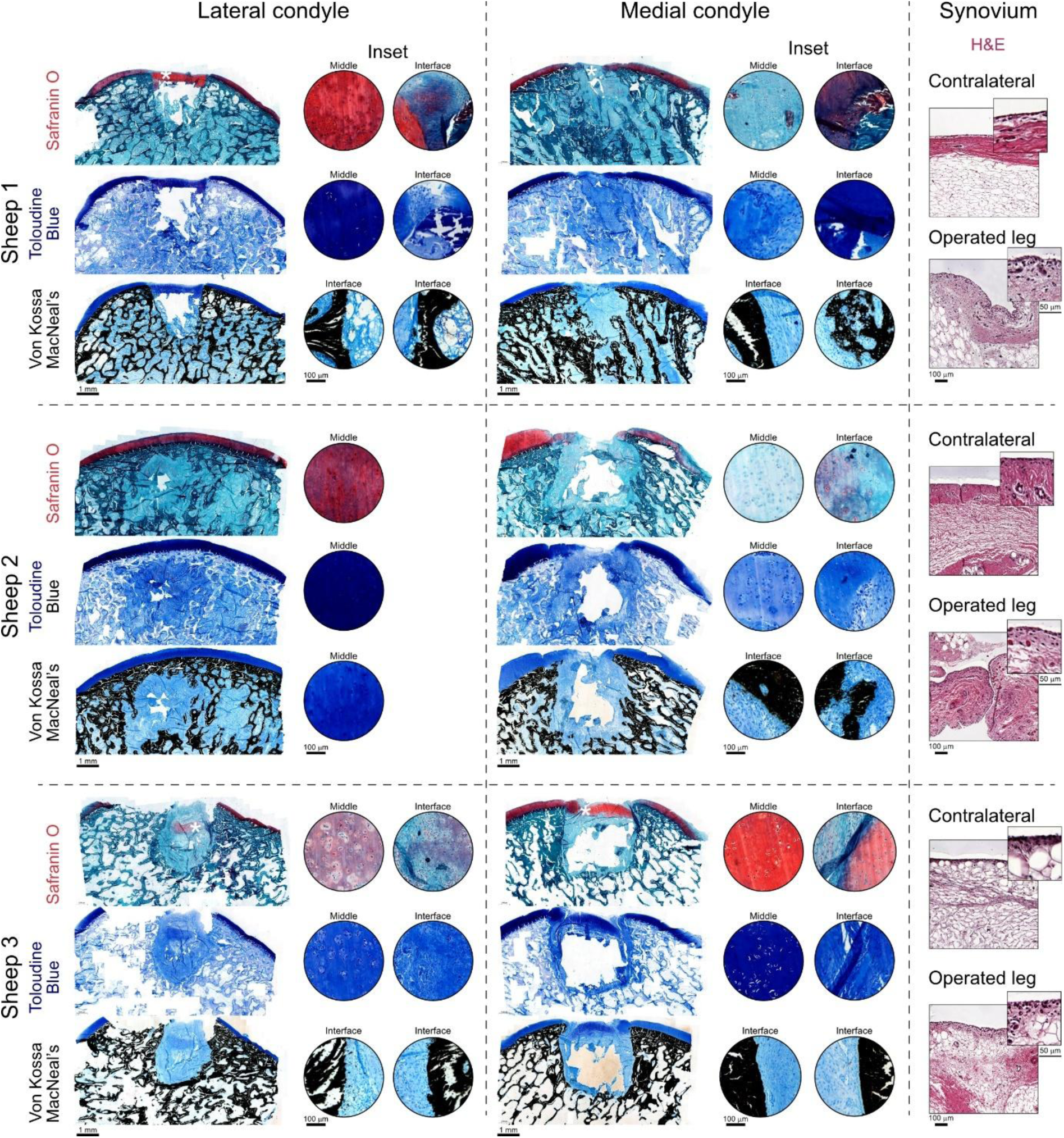
Evaluation of cartilage and bone infiltration on the osteochondral grafts (OCGs) implanted in an ovine model. Histological staining of 5 µm thin sections from explants (Sheep 1–3) in the lateral and medial condyles. Sections are stained with Safranin O, Toluidine Blue, and Von Kossa–MacNeal, with corresponding higher-magnification insets (within the engineered cartilage region and at the interface between the implant and the host-tissue). The location of the engineered cartilage is indicated by a white asterisk in the Safranin O images. Scale bar = 1 mm; inset = 100 µm. Synovial tissue from contralateral and operated joints is shown with H&E staining. Scale bar = 100 µm; inset = 50 µm.

The synovial membrane of both the operated joint and the healthy joint was collected to perform H&E staining and evaluate the degree of synovitis. Synovitis can be evaluated by the enlargement of the lining cell layer, cellular density of the synovial stroma, and leukocyte infiltration.^62^ In all cases, at the implantation site, the lining cells form a single layer, the synovial stroma shows normal cellularity, and there were no signs of an inflammatory infiltrate. Therefore, we suspect that there was no major inflammation in the joint cavity at the time of sample collection, 12 weeks post-implantation.

## Discussion

In this study, we demonstrated that functionally mature cartilage can be engineered from autologous nasal chondrocytes and that stable integration between cartilage and bone can be achieved through controlled hydrogel infiltration followed by cartilage maturation. Nasal chondrocyte-derived grafts reached instantaneous compressive moduli comparable to native sheep articular cartilage, whereas equilibrium modulus, dry weight, and glycosaminoglycan (GAG) content remained lower. Controlled infiltration of the hydrogel into the trabecular pores of the ceramic scaffold created a mechanical interlock that efficiently anchored the engineered cartilage to the bone compartment. Together, these findings demonstrate the feasibility of generating structurally integrated OC grafts containing mechanically mature cartilage. However, successful translation will require improved implant fixation within the OC defect, as inadequate press-fit likely led to osteolysis and implant sinking after implantation.

Applying our previously established hydrogel formulation and culture strategy for auricular chondrocytes^46^ to nasal chondrocytes resulted in cartilage with instantaneous compressive moduli comparable to native sheep articular cartilage. These values exceeded biomechanical moduli reported for OCG generated from equine articular chondroprogenitor cells (20 ± 6 kPa)^29^ and caprine microtissues composed of a 3:1 mixture of mesenchymal stromal cells and chondrocytes (262 ± 41 kPa)^27^, although differences in measurement should be acknowledged. They also exceeded the mechanical properties reported for engineered human nasal cartilage in a recent clinical study employing the nose-to-knee approach (approximately 10 to 100 kPa)^45^. Histologically, the engineered grafts closely resembled native hyaline cartilage through abundant GAG and collagen II deposition together with the absence of collagen I. Nevertheless, equilibrium modulus, dry weight, and GAG content remained below those of native cartilage. Given the close relationship between extracellular matrix composition and cartilage mechanics^63^, these findings suggest that although the grafts reached functional maturity, further extracellular matrix deposition and organization will be required to fully recapitulate native articular cartilage.

Stable integration between cartilage and bone was achieved by tuning the hydrogel’s viscosity to control infiltration into the trabecular pores of the ceramic scaffold. As the cartilage matured, the hydrogel pillars occupying these pores stiffened, progressively reinforcing the mechanical interlock between both compartments. This strategy resulted in an interfacial shear strength approaching those reported for the native OC interface^58^. Further strengthening of this transition zone may be achieved by introducing a calcified cartilage layer that more closely reproduces the native OC interface. Such an approach could promote localized calcification at the interface while preserving a hyaline cartilage phenotype at the articulating surface. Importantly, the robust cartilage maturation achieved with nasal chondrocytes allowed stable OC integration without requiring reinforcement of the cartilage layer.

Despite generating mechanically competent cartilage and a stable cartilage-bone interface, successful OC repair remained limited by implant fixation and subchondral bone integration. Previous studies using Sr-HT-Gahnite scaffolds for large bone defect repair have similarly demonstrated that scaffold fixation critically influences subsequent bone regeneration^53^. Although not performed in OC defects, it emphasizes the importance of stable mechanical fixation for successful osseointegration. Likewise, recent studies in equine^26^ and caprine^27^ OC defect models reported implant sinking following collapse of the surrounding bone, identifying insufficient subchondral support as a major failure point. Other groups have focused on reinforcing the cartilage compartment to prevent shear failure and compressive deformation^26^. Although such reinforcement substantially increased cartilage stiffness compared to non-reinforced hydrogels, the reported compressive moduli remained below those achieved by our nasal cartilage grafts, and implant sinking was still observed following implantation. Collectively, these findings suggest that fixation of the bone compartment in the surrounding bone becomes the dominant determinant of OCG performance, once cartilage reaches a certain maturity and mechanical stability. In our study, imperfect press-fit likely resulted in micromotion under repetitive joint loading, leading to osteolysis and allowing synovial fluid infiltration at the implant-bone interface, promoting bone cyst formation and ultimately implant sinking. Notably, sheep 1, which showed the most favorable integration and minimal osteolysis, also repeatedly removed its cast and was therefore exposed to earlier joint loading than sheep 2 and 3. While the tighter press-fit achieved in this animal is the most likely explanation for its superior outcome, we cannot exclude a contribution from earlier mechanical loading, which has been reported to promote osteochondral and bone graft integration in other models^64^.

In contrast to tissue-engineered OCG, autologous and allogeneic OCG transplantation has demonstrated reliable clinical integration with limited bone cyst formation or graft degeneration.^65–67^ A major advantage of these grafts is that they can be inserted under press-fit using tools such as the OATS instrumentation, thereby minimizing implant micromotion immediately after implantation. Although donor-site morbidity, limited graft availability, and potential disease transmission remain important drawbacks, tissue-engineered OCG insertion may be improved by developing specialized tools following these principles. Future scaffold designs could prioritize controlled radial compressibility of the bone compartment to facilitate reliable press-fit fixation while maintaining sufficient mechanical stability. At the same time, the mechanical stability of the engineered cartilage during implantation must be carefully considered to ensure that insertion forces required for stable implantation do not compromise cartilage integrity.

Several limitations should be acknowledged. First, this proof-of-concept study included only three animals and did not incorporate an untreated or clinical control group. Consequently, the study was designed to identify the principal failure mechanism rather than evaluate therapeutic efficacy. Second, the three-month follow-up primarily captures early healing events, and longer-term studies of six to twelve months will be required to assess durable OC regeneration. Additionally, implant micromotion was inferred from the observed osteolysis and implant sinking but was not quantified directly. Future studies integrating biomechanical measurements of primary implant stability will therefore be important to validate the proposed failure mechanism. Finally, fixation quality and joint loading exposure as observed in sheep 1, do not allow a definitive attribution of the favorable outcome to either factor alone and studies designed to test these factors individually and in combination are required.

## Conclusion

Taken together, our proof-of-concept study demonstrates that autologous, engineered OCG can be safely implanted, did not show biocompatibility issues and maintain cartilage-specific features in vivo. However, outcomes are highly dependent on achieving stable press-fit fixation. Insufficient implant–bone matching led to micromotion, osteolysis, and implant sinking within the first three months after implantation, highlighting the critical importance of precise defect matching and mechanical anchoring for successful OC regeneration. These findings highlight both the translational potential of the approach and the need for further optimization of implant design and surgical implantation technique to ensure reliable integration and long-term functionality.

## Experimental Section and Methods

### Nasal Cartilage Collection and Cell Isolation

The in vivo ovine experiments were conducted at the Musculoskeletal Research Unit according to Swiss laws for animal welfare and approved by the local governmental authorities (Kantonales Veterinäramt Zürich, Switzerland, licence no. ZH185/2022). Three female, skeletally mature Swiss Alpine sheep, (32 months of age, body weight average 61kg, range 59-63kg), from the Musculoskeletal Research Unit’s (MSRU) own herd were included in this study. Sheep were acclimatized under study conditions for at least seven days prior to the first surgery.

Sheep were fasted for a maximum of 24 hours with water available *ad libitum*. A pre-anesthetic examination was performed, including general clinical examination, hematology and chemistry panel. Thirty to forty-five minutes before induction, animals were premedicated intramuscularly (i.m.) with buprenorphine (0.01 mg/kg, Temgesic®, Reckitt Benckiser AG, Wallisellen, Switzerland) and medetomidine (0.005–0.01 mg/kg, Medetor®, Virbac AG, Opfikon, Switzerland). A pre-emptive dose of carprofen (4 mg/kg, Rimadyl®, Zoetis Schweiz GmbH, Zurich, Switzerland) was administered intravenously (i.v.) through a venous catheter. A booster for tetanus (3,000 IU/sheep, Tetanus Serum Intervet, MSD Animal Health GmbH, Luzern, Switzerland) was administered subcutaneously. Prophylactic antibiotics were administered intravenously (penicillin 30,000 IU/kg, Penicillin natrium Streuli ad us. vet., Streuli Pharma AG, Uznach, Switzerland; gentamicin 4 mg/kg, Vetagent® ad us. vet., MSD Animal Health GmbH, Luzern, Switzerland). Anaesthesia was induced intravenously with midazolam (0.1 mg/kg, Midazolam Sintetica, Sintetica AG, Mendrisio, Switzerland), ketamine (3–5 mg/kg, Ketanarkon® 100 ad us. vet., Streuli Pharma AG, Uznach, Switzerland) and propofol (0.4–4 mg/kg, Propofol® 1% Fresenius, Fresenius Kabi (Schweiz) AG, Kriens, Switzerland), administered to effect. Following laryngeal desensitisation with 10% lidocaine spray (Xylocain® Spray 10%, Aspen Pharma Schweiz GmbH, Baar, Switzerland), the trachea was intubated and correct placement was confirmed by expired carbon dioxide monitoring (EtCO₂). Anaesthesia was maintained with isoflurane (1–3%) in oxygen and air (FiO₂ 50–70%) delivered via an adult F-circuit. Monitoring included electrocardiogram, heart rate, pulse rate, non-invasive blood pressure, inspired and expired concentrations of carbon dioxide, oxygen and isoflurane, oesophageal temperature, and arterial oxygen saturation (SpO₂); parameters were recorded at 10-minute intervals.

Sheep were positioned in lateral recumbency with the head slightly inclined downwards. The nasal cavity was flushed with chlorhexidine and crusts were removed with a sterile sponge. Local anaesthesia of the nasal mucosa was achieved with 10% lidocaine spray immediately prior to incision. The mucosa was then incised and a full-thickness cartilage biopsy was harvested using a 6 mm diameter biopsy punch. Bleeding was controlled with electrocautery as required. The mucosa was closed with resorbable suture material (Vicryl 3/0, Ethicon, Johnson & Johnson, Spreitenbach, Switzerland). Post-operative analgesia consisted of buprenorphine (0.01–0.02 mg/kg i.m., every 4–6 hours on the day of surgery and carprofen (4 mg/kg s.c., once daily for 4 days). Antibiotic prophylaxis was continued for 4 days post-operatively.

Nasal cartilage from sheep were collected, washed in PBS containing 50 μg/mL gentamicin, freed from surrounding tissue and minced into 1 mm stripes. Cartilage was then digested in collagenase solution (DMEM 31966, 2% v/v FBS,1x Antibiotic-Antimycotic, 1000 CDU/mL collagenase from Clostridium histolyticum (C6885, Merck, CDU: collagen digesting units)) at 37°C for 4 h. Cells were collected by passing the obtained solution through a 40 μm cell strainer, centrifuging the solution at 250 rcf for 10 min, and collecting the cell pellet. Passaging from passage 0 to 1 was carried out by seeding 10’000 cells/cm^2^ in DMEM containing 10% (v/v) FBS, 50 μg/mL L-ascorbate-2-phosphate (TCI), 1x Antibiotic-Antimycotic, and 5 ng/mL FGF-2 (Recombinant Human FGF-basic, PeproTech). Expansion was carried out at 95% humidity, 5% CO_2_ and 37°C.

### Cell Culture

From passage 1 onwards, cells were cultured by seeding them at 3’000 cells/cm^2^ in DMEM containing 10% (v/v) FBS, 10 ng/mL FGF-2 and 10 ng/mL TGF-β3 (Animal Free Recombinant Human TGF-β3, PeproTech). At confluency (∼7 d), cells were trypsinized, centrifuged, the cell pellet collected, stained with Acridine Orange/Propidium Iodide Stain (F23001, Logos Biosystems) and cells counted on an automated fluorescence cell counter (LUNA-FX7, Logos Biosystems). Upon reaching passage 3, cells were trypsinized, counted and washed in TBG to remove any remaining calcium ions which could initiate crosslinking of the hydrogel. After washing, the cells were combined with the hydrogel and casted as detailed in the methods section “Hydrogel Casting”.

### In vivo Implantation and in-life Period

Fasting, pre-anaesthetic examination, premedication, intravenous catheterisation, induction and intubation were performed as described for the first surgery (nasal cartilage collection). Prophylactic antibiotics were administered as described for the first surgery.

Anaesthesia was maintained with a balanced protocol combining isoflurane (1–3%) in oxygen and air (FiO₂ 50–70%, delivered via an adult F-circuit) with variable-rate infusions of propofol (0.5–1 mg/kg/h) and ketamine (1.2–3 mg/kg/h). Ringer’s lactate solution was administered intraoperatively at 5–10 mL/kg/h. Monitoring parameters were as described for the first surgery, with the addition of invasive arterial blood pressure measurement (systolic, mean and diastolic) via a catheter placed in an auricular artery. Sheep were maintained under a warming blanket (Bair Hugger) throughout the procedure.

Following intubation, sheep were positioned in sternal recumbency for epidural analgesia. Preservative-free morphine (0.1 mg/kg, Morphin HCl Sintetica, Sintetica AG, Mendrisio, Switzerland), diluted in sterile 0.9% NaCl to a total injection volume of 0.05 mL/kg, was administered into the lumbosacral epidural space. At least ten minutes before surgical incision, loco-regional anaesthesia of the sciatic and femoral nerves of the operated hindlimb was performed under ultrasound guidance. The sciatic nerve was located at the midpoint between the major trochanter and the sciatic tuberosity, and the femoral nerve was identified in the connective tissue between the iliopsoas, pectineus and sartorius muscles at the level of the upper thigh. A volume of 0.1 mL/kg of ropivacaine 0.5% (Ropivacain Kabi, Fresenius Kabi (Schweiz) AG, Kriens, Switzerland) was injected at each site.

For the implantation surgery, sheep were positioned in dorsolateral recumbency with one hindlimb flexed at approximately 90° to the surgical table. The operated hindlimb was alternated between left and right from one animal to the next. The surgical site was clipped, scrubbed and draped according to standard aseptic protocol. An S-shaped skin incision of approximately 15–18 cm was made over the stifle joint, extending from the proximal patella medially to the level of the tibial tuberosity laterally. Soft tissue and fascia were incised parallel to the patellar ligament, and the medial and lateral retinacula were exposed.

A parapatellar approach was performed with the stifle joint in flexion to expose the weight-bearing areas of the medial and lateral femoral condyles using wound retractors. One cylindrical osteochondral defect was created in each condyle (two defects per animal). The defect site was first marked with a customized 6 mm metal biopsy punch positioned with a customized polymer drill guide to prevent slippage and damage to the adjacent cartilage. With the drill guide held in place, the punch was removed and a customized drill with a depth stopper was inserted to prepare the cylindrical graft bed. The drill diameter and depth were set per defect to match the corresponding implant, with a controlled press-fit clearance to account for the frictional properties of the ceramic. Depth was further adjusted to the local cartilage thickness, so that the upper surface of the engineered cartilage layer would sit level with the surrounding native articular cartilage. The resulting graft bed dimensions were 6.3 ± 0.1 mm in diameter and 6.4 ± 0.5 mm in depth (mean ± SD, n = 6, Table 1).

**Table 1.** Defect (drillhole) and OCG graft dimensions.

|  |  | Drillhole |  | OCG graft |  |
| --- | --- | --- | --- | --- | --- |
|  |  | Diameter [mm] | Depth [mm] | Diameter [mm] | Height (mm) |
| Sheep 1 | Lateral | 6.2 | 6.2 | 6.1 | 6.1 |
|  | Medial | 6.3 | 6.1 | 6.2 | 6.0 |
| Sheep 2 | Lateral | 6.3 | 6.1 | 6.1 (top)<br>6.2 (bottom) | 6.1<br>(overshot cartilage surface by 1.1mm) |
|  | **Medial** | 6.4 | 7.2 | 6.0 (top)<br>6.3 (bottom) | 7.1 |
| **Sheep 3** | **Lateral** | 6.2 | 6.2 | 6.1 | 6.1 |
|  | **Medial** | 6.3 | 6.7 | 6.1 | 6.4 |

After flushing the defect, the corresponding osteochondral implant was inserted into each defect using a customized applicator. Osteochondral implants were positioned so that the upper surface of the engineered cartilage layer was level with the surrounding native articular cartilage surface. Each finished osteochondral implant was individually measured prior to implantation (diameter 6.2 ± 0.1 mm; total height 6.3 ± 0.4 mm; mean ± SD, n = 6, Table 1).

The joint capsule was closed with resorbable suture material (Vicryl 2/0) and the skin was closed routinely with non-resorbable suture material (Supramid 2/0, B. Braun Medical AG, Sempach, Switzerland). Post-operative analgesia consisted of buprenorphine (0.01–0.02 mg/kg i.m., every 4–6 hours on the day of surgery) and carprofen (4 mg/kg s.c., once daily for 5 days).

Operated sheep hindlimbs were placed in a cast for a total of 5 weeks post-operatively to limit joint flexion. The cast was exchanged 3 weeks after surgery and skin sutures were removed. Sheep 1 however did not accept the cast and therefore its operated hindlimb was left without a cast after the first day following surgery. Sheep 2 accepted its cast for 4 weeks. Sheep 3 kept its cast for the planned duration. For the first three weeks after surgery, sheep were also placed in specialized suspension nets that would allow partial weight bearing but prevent uncontrolled movements. During this time period, sheep were housed in individual stalls, with eye and nose contact possible with their stall neighbors. Following cast removal, animals were housed in pairs, with a non-experimental companion sheep introduced to ensure none of the study sheep animals was housed alone. From six weeks post-operatively all animals were housed as a group, with access to indoor and outdoor space, until the end of the in-life phase, which was 12 weeks post-implantation.

Animals were monitored at least twice daily throughout the in-life phase by veterinarians, veterinary technicians or trained animal caretakers using standardised observation protocols. Appetite, posture, alertness, respiration, weight-bearing, gait and signs of pain were recorded using a semi-quantitative scoresheet, as well as any clinical abnormalities.

### Post-mortem macroscopic evaluation of implant sites

Twelve weeks after the implantation surgery, sheep were transported to the slaughterhouse of the Tierspital and sacrificed by captive bolt stunning followed by exsanguination, according to Swiss animal welfare law. The operated hindlimbs were harvested immediately. A layer-by-layer dissection of the operated stifle joints was performed and the femoral condyles exposed. A semi-quantitative macroscopic score was used to assess joint abnormalities, including the joint capsule, synovial tissue and liquid as well as the surrounding cartilage at the femoral condyles. Each implantation site was macroscopically assessed evaluating the cartilage repair according to ICRS guidelines, specifically looking at the degree of defect repair (percentage of filling height compared to surrounding cartilage), the integration to the border zone and the macroscopic appearance of the repair. Findings were also documented photographically.

### Histology – In vivo sample processing and assessment

Osteochondral blocks containing the defect, surrounding cartilage and underlying subchondral bone were prepared from each operated condyle and fixed in 40% ethanol to preserve the fluorochrome signal. Following fixation, samples were dehydrated through an ascending ethanol series (50% to 100%), cleared in xylene, and infiltrated under vacuum with methyl methacrylate (MMA) (Sigma-Aldrich Chemie GmbH, Steinheim, Germany). Polymerization was carried out at 4 °C, followed by sequential incubation at room temperature then at 37 °C until complete polymerization was achieved.

Polymerized blocks were cut along an anterior-posterior plane transecting the middle of the defect using a diamond saw (EXAKT 310, EXAKT Apparatebau GmbH & Co. KG, Norderstedt, Germany). One half of each block was used to produce thick ground sections (400–600 µm), which were mounted on white opaque acrylic slides (Perspex GS Acrylglas Opal, Wachendorff Kunststofftechnik, Basel, Switzerland) and used for surface-stained with Toluidine Blue. The other half of each block was trimmed and used to prepare thin sections (5 µm) on a microtome (Leica RM 2155, Leica Microsystems, Wetzlar, Germany), which were stained with Toluidine Blue, Safranin O / Fast Green, and von Kossa / McNeal.

Surface-stained sections were imaged using a Leica Z6 APOA microscope equipped with a Leica DFC 420C camera and image acquisition was done using the Imagic IMS software (Imagic Bildverarbeitung AG, Opfikon, Switzerland). Thin sections were imaged on an automated slide scanner (Panoramic 250, 3D Histech).

### LCD 3D printing of Sr-HT-Gahnite

Sr-HT-Gahnite scaffolds were fabricated using a ceramic vat-photopolymerisation approach adapted from our previously reported LCD-based bioceramic printing method^54^. The ceramic precursor consisted of Sr-HT powder blended with 15 wt% alumina powder, with a median particle size of approximately 4 µm, which was dispersed in a photocurable resin to form a ceramic-resin slurry. Briefly, the ceramic slurry was prepared at 40 vol% solid loading. Sodium polyacrylate was added as a dispersant at 1.5 wt% relative to the dry ceramic powder to improve particle dispersion and minimise sedimentation during printing. The liquid resin phase (Phrozen Aqua Clear) consisted of acrylate oligomer, 4-(1-oxo-2-propenyl)-morpholine, and Bis(1,2,2,6,6-pentamethyl-4-piperidinyl) sebacate. The ceramic powder was gradually introduced into the liquid phase under mechanical mixing to avoid agglomeration. The suspension was then transferred into milling jars containing zirconia milling media and homogenised by ball milling at 110 rpm for 8 h. Following homogenisation, the slurry was passed through a stainless-steel sieve (63 µm) to remove large agglomerates and obtain a uniform printable ceramic-resin suspension.

Trabecular scaffold geometries were generated from µCT scans of ovine subchondral trabecular bone. The trabecular region beneath the subchondral plate was segmented, converted into an STL file, and modified to improve printability while preserving the native trabecular architecture. Shrinkage compensation was applied to the digital models before printing to account for dimensional changes during debinding and sintering.

Scaffolds were printed using a LCD-based vat-photopolymerisation system equipped with a 7.1-inch projection LED display operating at 460 nm, with an in-plane pixel resolution of 22 µm. Prior to printing, silanised borosilicate glass slides were attached to the build platform to improve adhesion of the green ceramic constructs. Printing was performed using a layer thickness of 50 µm. The exposure time was optimised to ensure sufficient curing of each ceramic-resin layer while minimising over-curing and pore closure. Two initial bottom layers were printed using extended exposure to enhance attachment to the build platform. After printing, green scaffolds were washed in ethanol to remove uncured ceramic resin from the scaffold surface and internal pores. Residual slurry within the trabecular architecture was removed using repeated solvent washing and compressed air until the pores were visually confirmed to be open. The printed constructs were then carefully detached from the glass substrate using an ultrasonic cutter to minimise damage to the fragile green bodies. The cleaned green scaffolds were dried and subjected to a multistage heat-treatment schedule to remove the organic resin components and sinter the ceramic phase at 1250°C for 3h. The thermal cycle included slow heating through the debinding temperature range to prevent cracking from rapid polymer burnout, followed by high-temperature sintering to consolidate the Sr-HT-Gahnite ceramic structure. After sintering, scaffolds were allowed to cool to room temperature inside the furnace. The sintered ceramic scaffolds were characterised by µCT and SEM to confirm preservation of the trabecular architecture, pore interconnectivity, surface morphology, and dimensional fidelity. Prior to hydrogel casting and osteochondral graft fabrication, scaffolds were sterilised by autoclaving and handled aseptically.

### Material Synthesis

HATG was synthesized as reported previously.^47,68^ Briefly, hyaluronan (HA, Lifecore Biomedical) was thiolated with 3,3’-dithiobis(propanoic dihydrazide) followed by vinyl sulfonation through the addition of divinyl sulfone. In a last step, glutamine (NQEQVSPL-ERCG) or lysine (FKGG-ERCG) donor peptides were allowed to react with the vinyl sulfonated hyaluronan. The product was sterilized by filtering through a 0.2 μm filter and lyophilized. A degree of substitution of ∼10-13% over the disaccharide repeating units was determined by nuclear magnetic resonance (NMR) and crosslinking dynamics confirmed by rheology.

### Scanning electron microscopy (SEM)

Ceramic samples were attached to metallic stubs using conductive carbon paste. Prior to imaging, surfaces were coated with a 10 nm carbon layer using a CCU-010 Carbon Coater (Safematic). Morphological analysis was carried out using a Zeiss Merlin SEM at an accelerating voltage of 2 kV.

### Hydrogel Casting

HA, HATG and Alg were dissolved in TRIS-buffered glucose (TBG, 50 mM TRIS, 200 mM d-glucose, pH 7.6). To prepare hydrogels, the HATG-Alg-HA solution was combined with factor XIII (FXIII, stock: 200 U/mL, Fibrogammin, CSL Behring), thrombin (stock: 500 U/mL, TISSEEL, Baxter) and the cell pellet to reach final concentrations of 0.5% (w/v) HATG, 0.25% (w/v) Alg, 0.75% (w/v) HA, 10 U/mL FXIII, 12.5 U/mL thrombin and 30 million cells per mL. Gels were then cast into sterilized PDMS molds either with or without the bone ceramic, sprayed with 100 mM CaCl2 to initiate crosslinking using an atomizer bottle and incubated for 15 min. Afterwards, 2 mL of 100 mM CaCl2 were added and incubated for another 45 min to finalize crosslinking. Once crosslinked, samples were washed two times in TBG to remove excess CaCl2 and cultured in chondrogenic media (DMEM 31966 supplemented with 50 μg/mL L-ascorbate-2-phosphate, 40 μg/mL L-proline, 10 μg/mL gentamicin, 1% (v/v) ITS+ (ITS+ Premix Universal Culture Supplement, Corning) and 10 ng/mL TGF-β3 – control discs: 2 mL, OCGs: 3 mL).

### Rheology

Rheological characterization was carried out on an MCR302e rheometer (Anton Paar) equipped with a Peltier element (P-PTD 200, Anton Paar) and thermal hood (H-PTD 200, Anton Paar). All tests were carried out at 25°C and a wet tissue placed around the samples to prevent dehydration. A parallel plate geometry with a 20 mm diameter was used (PP-20, Anton Paar) and a gap of 200 μm set.

Crosslinking tests (time sweeps) were carried out at a frequency of 5 rad/s, oscillatory shear strain of 1% and measurement point duration of 10 s. Samples were crosslinked by adding 10 μL of 1M CaCl_2_ around the sample to initiate crosslinking and simulate the CaCl_2_ spraying used during sample preparation in cell culture. After 1 h, 500 μL of 100 mM CaCl_2_ were added. Samples were allowed to crosslink for 4 h. After crosslinking, stress-relaxation tests were carried out by applying a step strain of 15% and measuring sample relaxation for 6 h, with measurement points recorded every 10 s. The relaxation half time was calculated as the time it took the sample to relax to half of its initial minus final value.

Shear thinning tests were carried out in rotational tests by logarithmically increasing the shear rate from 0.01 s^-1^ to 300 s^-1^ while the data point duration was logarithmically decreased from 300 s to 1 s.

### Microscopy

Viability imaging was performed by staining cells in DMEM 31966 supplemented with 1 μM Calcein AM (Invitrogen) and 1 μM propidium iodide (PI, Fluka) for 1 h. Afterwards, samples were briefly washed in DMEM 31966 and imaged on a multiphoton microscope (Leica SP8 MP) equipped with a 25x water immersion objective. Calcein AM and PI excitation was carried out at 900 nm and their emission collected at 500–570 nm and 695–740 nm respectively. Second harmonic generation was collected simultaneously at half the excitation wavelength (442–458 nm) and confirmed as sharp peak by moving the slider outside the measurement window and observing signal disappearance. Samples were imaged from the sample surface 100 μm into the sample at 1μm z-steps. Viability was calculated by counting viable and dead cells at 0, 25, 50, 75 and 100 μm and dividing the number of viable cells by the total number of cells (viable and dead) over the entire range.

### Mechanical Characterization

Unconfined stress-relaxation compression testing was carried out on a TA.XTplus texture anlyzer (Stable Micro Systems) equipped with a 500 g load cell. Samples were imaged on a stereomicroscope (Leica Wild Heerbrugg) and the cross-sectional area of the sample determined in ImageJ2 (v2.16.0/1.54p). The samples’ surface was trimmed with a microtome blade to ensure a flat surface. Samples were placed between the compression plates and 200 uL of saline solution (0.9% NaCl) added to prevent dehydration of the sample. An initial pre-strain of 15% was applied to the sample to ensure full contact between the sample and the compression plates. Testing was carried out by applying four consecutive 5% strain intervals at a strain rate of 5%/s to reach a final strain of 20%. After the first two intervals, samples were allowed to relax for 30 min and after the last two intervals for 45 min. The instantaneous modulus was calculated by linear fitting the stress-strain curve of the last loading interval from 15-20% and the equilibrium modulus was calculated by linear fitting the corresponding stress-strain values at equilibrium at 0%, 5%, 10%, 15% and 20% strain.

Shear tests were performed using a custom mount allowing to clamp the bone ceramic part of the OCG and shear the cartilage part off along the radial direction. A custom shear interface was lasercut in a half-moon shape and a 6 mm diameter to allow direct contact with the cartilage’s radial surface. The cartilage was sheared at 0.5 mm/s. The shear stress was calculated by using the cross-sectional area of the OCG and the shear stress at breakage reported as the highest shear stress during the test.

### Histology – Paraffin Sections

Samples were fixed in 4% paraformaldehyde for 4 h and subsequently dehydrated in graded ethanol solutions. Paraffin embedding was performed on an automated tissue processor (Milestone LogosJ) and 5 μm sections cut on a microtome. Prior staining samples were deparaffinized in xylene and rehydrated. Stained sections were imaged on an automated slide scanner (Panoramic 250, 3D Histech).

#### Safranin O

Sections were stained in Weigert’s iron hematoxylin solution (HIT107, Merck) for 5 min, washed in deionized water, differentiated in 1% acid alcohol (1% of 37% HCl in 70% ethanol) for 2 s, washed in deionized water, stained in 0.02% fast green (F7252, Merck) for 1 min, differentiated in 1% acetic acid for 30 s, stained in 1% safranin O (S8884, Merck) for 30 min, dehydrated to xylene and mounted.

#### Alizarin Red

Sections were stained in 2% alizarin red solution (A5533, Merck) for 2 min, washed in deionized water, dehydrated to xylene and mounted.

#### Picrosirius Red

Sections were first stained in Weigert’s iron hematoxylin solution (HIT107, Merck) for 5 min, followed by washing in deionized water. Slides were then incubated for 1 h in 0.1% of Direct Red 80 in a saturated solution of picric acid. Finally, slides were washed in acidified water, dehydrated to xylene, and mounted.

#### Elastin

Sections were stained using the Elastic Stain kit according to the manufacturer’s instructions (HT25, Merck). Briefly, slides were stained in working elastic stain solution for 10 minutes, differentiated in working ferric chloride solution, counterstained in Van Gieson Solution for 2 minutes, dehydrated to xylene and mounted.

#### Immunohistochemistry

Antigen retrieval was carried using hyaluronidase (8000 U/mL in PBS, H3506, Merck) at 37°C for 30 min, sections subsequently washed in PBS, blocked with 5% BSA in PBS at room temperature for 1 h and incubated with the primary antibody (collagen I: mouse anti-collagen I, 1:1000, stock: 1.3 mg/mL, ab6308, Abcam; collagen II: mouse anti-collagen II, 1:20, stock: 75 μg/mL, Hybridoma Product II-II6B3, deposited by Linsenmayer, T.F.^69^; collagen X: mouse anti-collagen X, 1:400, stock: 0.5 mg/mL, 14-9771-80, eBioscience) in 1% BSA in PBS at 4°C overnight. The next day, slides were washed in PBS, endogenous peroxidase quenched in 0.3% H_2_O_2_ for 15 min, sections washed again in PBS and incubated with the secondary antibody (goat anti-mouse IgG-HRP, 1:1000, stock: 2 mg/mL, ab6789, Abcam) in 1% BSA in PBS for 1 h at room temperature. Sections were washed in PBS and developed in chromogen solution (DAB substrate kit, ab64238, Abcam). Afterwards, sections were washed in deionized water, stained in Weigert’s iron hematoxylin solution (HIT107, Merck) for 3 min, washed in deionized water, destained in 1% acid alcohol (1% of 37% HCl in 70% ethanol) for 2 s, washed, blued in 0.1% Na_2_CO_3_ for 1 min, washed, dehydrated to xylene and mounted.

#### H&E

Sections were first stained in hematoxylin solution modified according to Gill III (1051740500, Merck) for 5 min, washed under running tap water for 10 min, differentiated in 95% ethanol for 30 s, followed by bluing in 0.1% Na_2_CO_3_ in deionized water for 40 s and dipped 10 times in 95% ethanol. The sections were then submerged in 0.25% Eosin Y (230251, Merck) prepared in 80% ethanol with 0.5% acetic acid for 1 min, washed 3 times in 100% ethanol, dehydrated in xylene and mounted.

### ECM Quantification

#### Dry Weight

To determine the dry weight of grafts and control cartilages, samples were weighted, placed in Eppendorf tubes, lyophilized for 48 h and weighted again. The dry weight percentage was calculated as weight after lyophilization (dry weight) divided by the weight before lyophilization (wet weight).

#### Sample preparation

To prepare the engineered cartilage and control cartilages for characterization, a papain solution was made by combining 10 mM L-cysteine HCl (C-7477, Sigma), 100 mM sodium phosphate (567547, Merck), and 10 mM Ethylenediaminetetraacetic acid (EDTA, A2937, AppliChem) in MilliQ water. To achieve a pH of 6.3, the solution was adjusted with 1 M HCl. Papain at 270 µg ml^−1^ was incorporated immediately before use in the digestion solution. Then, 500 µl were added to approximately 10 mg of wet weight sample material (exact quantities were recorded, and concentrations were calculated accordingly). The solutions were digested at 60 °C overnight with shaking at 1000 rpm.

#### DNA

The amount of DNA was quantified using a Quant-iT PicoGreen dsDNA assay kit (Invitrogen) following the manufacturer’s protocol. Briefly, 1X Tris-EDTA (TE) Buffer was prepared by diluting 1:20 the 20X TE buffer from the kit in MilliQ-water. The PicoGreen was then diluted 1:200 in the 1X TE buffer. DNA standard solutions with final concentrations of 1000, 500, 100, 10, and 0 ng ml−1 were prepared by diluting the 100 µg ml^−1^ DNA standard from the kit in 1x TE buffer. Then, 10 µl of the digested cartilage samples were distributed into a 96-well plate. These were each diluted with 40 µl of 1X TE buffer. The DNA standards were placed on the plate with 50 µl each. In all samples, 50 µl of PicoGreen solution was added, resulting in a final dilution of 1:10. Duplicates were performed for all samples. The fluorescence was measured with a microplate reader (Synergy H1 Hybrid Reader, BioTek) at 480 nm excitation and 520 nm emission. The resulting data were analyzed in Excel to obtain a standard curve, from which the DNA concentration in the tested samples was calculated.

#### GAGs

GAG content was quantified using the Blyscan sGAG assay according to the manufacturer’s instructions. First, GAG standards were prepared containing 0, 1, 2, 3, 4, and 5. µg of the reference standard and mixed with the digestion buffer (without papain). Test samples of cartilage and engineered cartilage were diluted 1:200, whereas the dECM was diluted 1:100. Then, the Blyscan dye reagent was added to each tube, and the tubes were placed on a mechanical shaker for 30 min at 100 rpm. During this time, a sulfated GAG-dye complex formed and precipitated out of the soluble, unbound dye. Later, the tubes were transferred to a centrifuge and spun down at 12 000 rpm for 10 min. The tubes were then inverted and drained by gently tapping them on a tissue. The dissociation reagent was added to each tube in a volume of 0.25 ml and vortexed to release the bound dye into solution. 100 µl of each sample were transferred to individual wells in a 96-well plate by non-rapid pipetting, to avoid foam formation, which could cause abnormal absorbance readings. The measurement was performed in a microplate reader (Synergy H1 Hybrid Reader, BioTek) with 656 nm absorbance. The resultant data was analyzed with Excel to obtain a standard curve, from which the concentration of GAG content in the tested samples was calculated.

### Micro-computed Tomography

Micro-computed tomography (μCT) was performed on a μCT 45 device (Scanco Medical, CH). μCT scans of the osteochondral unit to generate the trabecular bone 3D model were acquired at 90 kVp, 44 μA and 4 W using a 0.1 mm Cu filter (3 μm voxel size). The image data were recorded using a sampling time of 2’000 ms and 2000 projections/180°. After harvesting the joints, scans of the explants were acquired in 0.9% (w/v) NaCl, at 90 kVp, 88 μA and 8 W using a 0.1 mm Cu filter (14.6 μm voxel size). The image data were recorded using a sampling time of 1’500 ms, 2000 projections/180°. Automatic reconstruction was performed in the Scanco software and exported as DICOM files. 3D reconstructions were carried out in Mimics (v.25, Materialise, Belgium) and 3-matic (v.17, Materialise, Belgium).

### Large Language Models (LLMs)

Large language models (ChatGPT, OpenAI) were used for language editing and proofreading during the preparation of the manuscript. Accuracy and factual correctness were ensured by the authors. No scientific content or analysis was generated using these tools. Use of LLMs followed the “Guide for Users of AI at ETH Zurich” published on July 24, 2025.

### Statistical Analysis

All experiments were performed with three independent replicates (n = 3). Data are presented as mean ± standard deviation. Statistical analysis was performed using ordinary one-way analysis of variance (ANOVA) followed by multiple comparisons testing to evaluate differences between groups. Except for the shear test, where statistical significance was assessed using an unpaired t-test with Welch’s correction. Statistical significance was defined as p < 0.05. Exact p-values are indicated directly in the graphs.

## Acknowledgements

This work was supported by the Swiss National Science Foundation (grant number: 10000207) and by the Australian Research Council Discovery Project DP220102876. The in vivo experiment consumables were funded by an interdisciplinary grant received from the Vetsuisse Faculty at the University of Zürich. The authors gratefully acknowledge ScopeM for their support and assistance in this work. The authors further gratefully acknowledge Lucrezia Baldi for assisting with the sheep surgery and supporting graft culture as well as the veterinary and animal care team at the MSRU (Dr.med.vet. Peter Kronen Dipl. ECVAA, Dr.med.vet. Marta Kantyka, Dr.med.vet. Olivia d/Anselme Dipl. ECVAA, Dr.med.vet Katja Nuss, med.vet. Valerie Eckart, med.vet. Benedikt Schlusemed.vet. Christopher Zillinger, med.vet. Tina Claussner, Rosita Walther, and Joanna Schmid) who helped with the sheep study and the MSRU laboratory technician (Aymone Lenisa) who performed the histological processing of the in vivo study samples.

## Data Availability Statement

The data generated and analyzed in this study are available in the ETH Zurich Research Collection repository under the doi: 10.3929/ethz-c-000807281.

## Supporting information

**Figure S1.**
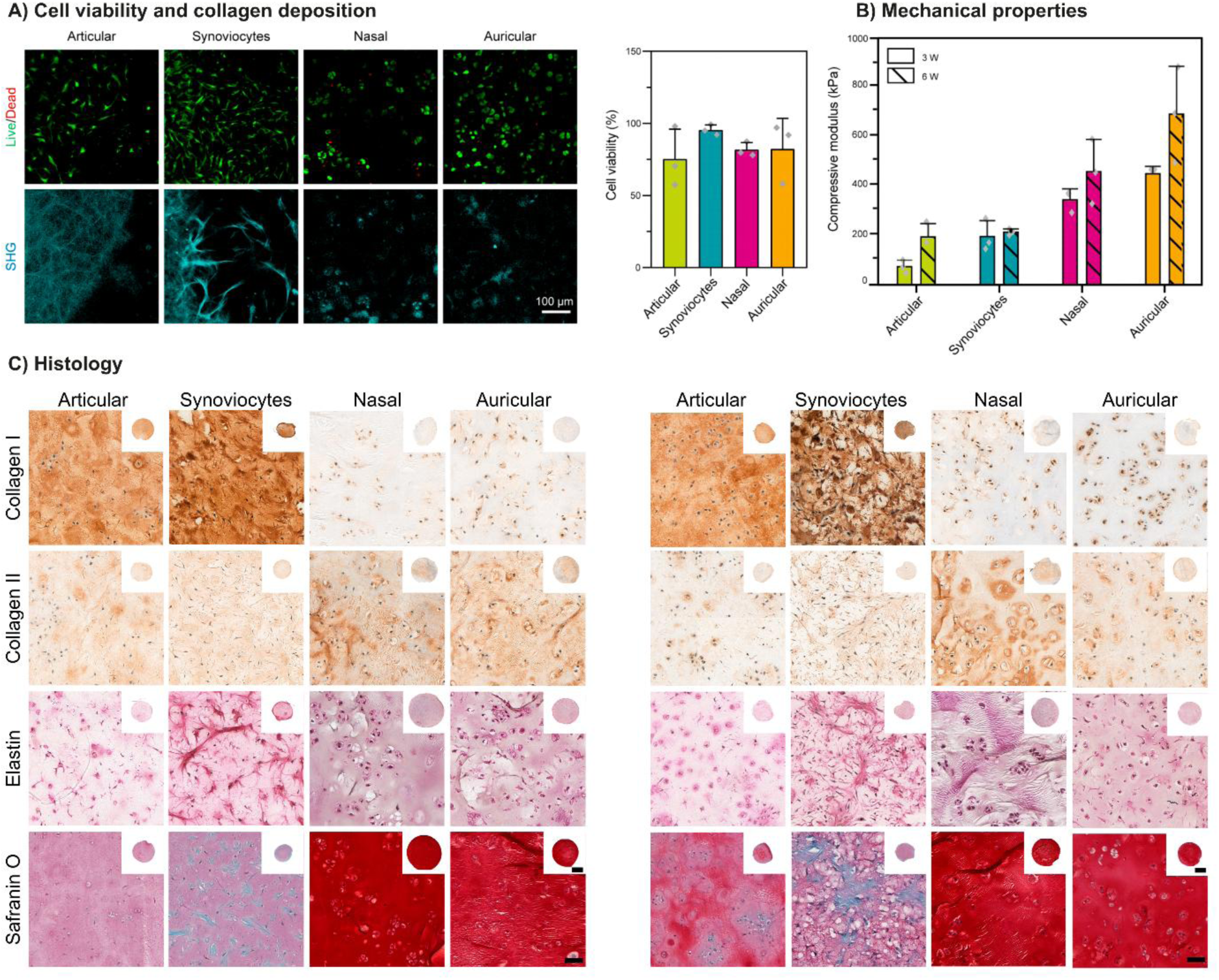
Cell source characterization. **(A)** Cell viability and matrix formation assessed by live/dead staining after 3 weeks (3W), and collagen deposition visualized by second harmonic generation (SHG) imaging. **(B)** Mechanical characterization showing the instantaneous modulus of the different groups at 3W and 6W. **(C)** Histological and immunohistochemical characterization of engineered grafts (6W), including Safranin O staining and elastin for histology, and collagen I, II immunohistochemistry. Scale bar = 50 µm; inset = 1mm.

